# A general approach to explore prokaryotic protein glycosylation reveals the unique surface layer modulation of an anammox bacterium

**DOI:** 10.1101/2020.12.03.409086

**Authors:** Martin Pabst, Denis Grouzdev, Christopher E. Lawson, Hugo B.C. Kleikamp, Carol de Ram, Rogier Louwen, Yuemei Lin, Sebastian Lücker, Mark C.M. van Loosdrecht, Michele Laureni

## Abstract

The enormous chemical diversity and strain variability of prokaryotic protein glycosylation makes a large-scale exploration exceptionally challenging. Therefore, despite the universal relevance of protein glycosylation across all domains of life, the understanding of their biological significance and the evolutionary forces shaping oligosaccharide structures remains highly limited.

Here, we report on a newly established mass binning glycoproteomics approach that establishes the chemical identity of the carbohydrate components and performs untargeted exploration of prokaryotic oligosaccharides from large-scale proteomics data directly. We demonstrate our approach by exploring an enrichment culture of the globally relevant anaerobic ammonium-oxidizing bacterium *Ca.* Kuenenia stuttgartiensis. By doing so we resolved a remarkable array of oligosaccharides, produced by two entirely unrelated glycosylation machineries targeting the same surface-layer protein (SLP) simultaneously. More intriguingly, the investigated strain also accomplished modulation of highly specialized sugars, supposedly in response to its energy metabolism—the anaerobic oxidation of ammonium —which depends on the acquisition of substrates of opposite charge. Ultimately, we provide a systematic approach for the compositional exploration of prokaryotic protein glycosylation, and reveal for the first time a remarkable balance between maximising cellular protection through a complex array of oligosaccharides and adhering to the requirements of the ‘metabolic lifestyle’.

---

Post-translational modifications modulate protein properties in response to the environment at very short time scales.^[1]^ Thereof, protein glycosylation is fundamental to all three domains of life and likely the most abundant modification out of the many reported to date.^[2]^ Moreover, glycoproteins are also found in protective envelopes of most pathogenic viruses.^[3]^ Carbohydrates, the building blocks of glycans, are the least conserved class of molecules and, when linked to proteins, considerably increase the proteome diversity. Consequently, due to glycosylation and the many other types of modifications, the number of proteoforms is orders of magnitude larger than what can be translated from the genomic sequence of an organism alone.^[4]^

Apart from well-investigated pathogens or easily culturable model archaea, such as *C. jejuni or H. volcanii*, our understanding of the biological significance and evolutionary forces shaping glycan structures is highly limited. This lack of understanding is largely due to the general paucity of large-scale approaches, capable of resolving the chemical diversity expressed by prokaryotes. In eukaryotes, protein glycosylation utilizes a well-defined number of some 10 monosaccharides. Thereby, the existence of distinct glycosylation systems and oligosaccharide structures evolved as ubiquitous for many molecular processes, such as protein folding, stability and immunity.^[4b, 5]^ In contrast, biosynthetic routes in prokaryotes have a much higher diversity.^[4b]^ Oligosaccharide chains show large variations in regard to sugar and linkage chemistry across species and strains.^[6]^ Interestingly, many functions protein glycosylation serves in eukaryotes, such as for protein folding, quality control or intracellular trafficking, do not apply to prokaryotes.^[7]^ However, glycan biosynthetic routes involve many enzymatic steps and are, particularly for prokaryotes, an energetically costly process. Therefore, those structures must serve other, but likely equally important roles.^[8]^

Prokaryotes account for a substantial proportion of Earth’s biomass, drive many global processes such as the nitrogen cycle or methane production, and directly impact human health (e.g., as exemplified by the gut microbiome).^[9]^ Anaerobic ammonium-oxidizing (anammox) bacteria, a phylogenetically deep branching group within the Planctomycetes phylum, play a key role in the bio-geochemical nitrogen cycle (e.g., producing over two thirds of atmospheric N_2_)^[10]^ and are the cornerstone of new, sustainable and resource-efficient biotechnologies (e.g., wastewater treatment).^[11]^ Members of the Planctomycetes phylum display cellular characteristics that are unique and strikingly homologous to those found in eukaryotes, such as cytosolic compartmentalization and endocytosis-like uptake of macromolecules and proteins.^[12]^ More specifically, the intra-cytoplasmic cellular compartment (anammoxosome), which is home to anammox catabolism, may be considered a functional analogue of the eukaryotic mitochondrion.^[12b]^ However, the recently confirmed existence of a peptidoglycan layer supported their classification as Gram-negative bacteria.^[13]^ Although so far only little is known about protein glycosylation in planctomycetal bacteria, pioneering work by van Teeseling et al. (2014 and 2016) revealed the existence of a glycosylated cell surface layer in planktonically growing *Ca.* Kuenenia stuttgartiensis. Moreover, a follow-up study by Boleij et al. (2018), reported on a glycosylated surface layer protein in the closely related genus *Ca.* Brocadia sapporoensis.^[14]^

Historically, protein glycosylation in prokaryotes has been associated with pathogens (and commensal bacteria), many of which display complex oligosaccharide structures, including highly specialized sugars such as nonulosonic acids (NulOs, frequently referred to as ‘sialic acids’).^[15]^ These sugars, in addition to cellular protection, have been shown to play key roles during pathogenicity and host invasion, such as in *C. jejuni,* which is one of the most common causes of food poisoning globally.^[16]^ Nevertheless, protein glycosylation is also common to non-pathogenic bacteria and archaea, commonly modulating surface layer proteins (SLPs), pili or flagella.^[6, 16c]^ This has been shown to introduce remarkable properties, such as for the SLP glycosylation of the extremophile *Sulfolobus islandicus* which provides the physical resistance to survive under extreme conditions.^[17]^ Nevertheless, most likely due to the lack of large-scale data, there is a general belief that glycosylation in (non-pathogenic) prokaryotes mediates structural, rather than functional, properties.^[18]^

From an analytical perspective, the enormous chemical and structural diversity expressed in prokaryotes makes large-scale exploration exceptionally challenging. High-performance (glycan database utilizing) approaches^[19]^ cannot be readily applied to prokaryotic glycoproteomes with unknown oligosaccharide chemistry. Therefore, the glycoproteomic exploration of novel prokaryotic strains often still employs protein fractionation and carbohydrate-specific colorimetric staining.^[15]^ The target proteins and associated carbohydrate structure(s) are then commonly identified following proteolytic digestion via mass spectrometric approaches. Thereby, peptides are sequenced and investigated for spectra sharing peptide and oligosaccharide fragments.^[14a, 14b, 15, 20]^ This approach takes advantage of the commonly observed monosaccharide fragments (oxonium ions) to differentiate unmodified peptides from glycopeptide spectra. However, this multi-step, manual procedure requires extensive experience and is extremely time consuming. Furthermore, the oxonium markers, supporting (automated) identification for eukaryotic proteins, are not necessarily part of the prokaryotic carbohydrate chains. Therefore, this strategy is heavily operator biased and may leave a large number of oligosaccharide modifications unidentified.

Recently, open modification search approaches, which match fragment ions to a protein sequence database without prior consideration of the intact peptide mass, have advanced the search for unexpected peptide modifications.^[21]^ Very recently, this approach has been combined with glycopeptide enrichment to analyze a range of pathogenic bacteria.^[22]^ Thereby, the enrichment improves spectral coverage, reduces the search space and qualifies observed mass-shifts as carbohydrate-like modifications. Moreover, an additional ion mobility interface (FAIMS) showed promising outcomes even without prior enrichment procedures.^[23]^ Those developments have significantly advanced the identification of oligosaccharide-related modifications in prokaryotic proteomes. However, until now, the large-scale discovery of prokaryotic protein glycosylation remains depended on specific enrichment procedures (by HILIC or specialized equipment), extensive fragmentation experiments (to ensure peptide sequence coverage) and proteome sequence database availability. The number of sequencing spectra and the database volume, furthermore, impact computational efforts and sensitivity. Nevertheless, common large-scale glycoproteomics approaches have been also established for the application to pure cultures. Most importantly, however, those approaches do not establish the strain-specific sugar components or additional oligosaccharide sequence information. Therefore, the untargeted large-scale exploration, which provides additional chemical and compositional information to support the physicochemical interpretation of the oligosaccharide chains—in particular, when exploring non-pure or enrichment cultures— remains a bottleneck in microbial proteome research.

We established a systematic procedure to analyze yet-unexplored prokaryotic protein glycosylation directly from large-scale proteomics data. By this means, we first determine the strain-(or enrichment)-specific sugar components and subsequently the corresponding protein linked oligosaccharide chains of individual strains in the culture (community). When investigating an enrichment culture of the globally relevant anammox bacteria, the new approach resolved a remarkably complex array of surface-layer oligosaccharides generated by two glycosylation machineries simultaneously. The physicochemical interpretation ultimately suggests an evolutionary link to charged surface layer proteins (SLPs) and the requirements of the metabolic lifestyle in anammox bacteria.

## RESULTS

### THE MICROBIAL GLYCOPROTEOMICS APPROACH

Establishing strain-specific monosaccharide components from proteomics data, the first step in our approach, takes advantage of the uniform fragment-ion patterns obtained when mass binning thousands of individual peptide sequencing spectra from shotgun proteomic data (Figure 1A–B, SI-DOC-B4). The mass patterns are highly comparable across species because they reflect the general amino acid repertoire used throughout all domains of life. The low mass range is dominated by mostly intense fragments of the canonical set of amino acids and combinations thereof (mono, di- and trimers). Carbohydrate-related signals, however, typically lay outside the amino acid fragment mass space due to their distinct chemical composition (SI-DOC-B1:3,5). Previously, mass binning had been proposed to search for stable amino acid modifications.^[24]^ Here, we explore the ‘out-of-range’ peaks from high-resolution data in search of (novel) carbohydrate-related signals. ‘Non-amino-acid-signals’ are then automatically accessed for their chemical composition at sub-ppm (parts per million) accuracy (Figure 1B:C). Thereby, candidates are analysed using a constructed sugar composition database, containing more than 3300 theoretical chemical sugar compositions, occurrence of additional water loss and co-occurrence within sequencing spectra (SI-DOC-B1:3). To provide maximum differentiation from closely related compositions of peptide fragments, this step employs shotgun proteomic data for which the fragmentation spectra were acquired at very high mass resolution (140K at m/z 200). Moreover, to provide in-depth chemical information from novel sugar derivatives, selected components (such as NulOs) were analyzed by additional in-source fragmentation experiments (SI-DOC E2).

**FIGURE 1.**
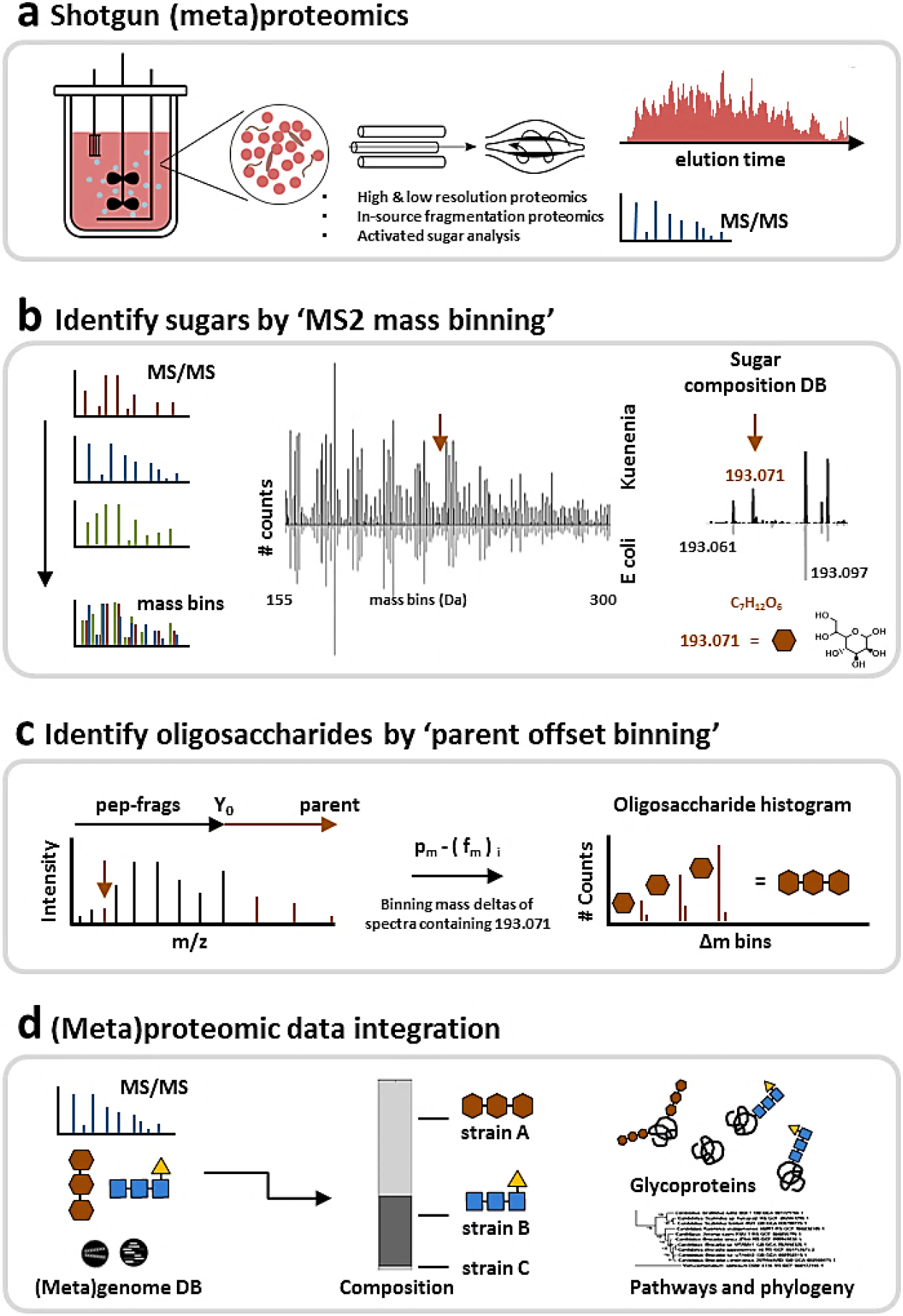
The mass binning glycoproteomics approach to explore prokaryotic protein glycosylation. **a)** The approach identifies the chemical composition of the proteome specific carbohydrate components and establishes the related protein linked oligosaccharide chains from large-scale (meta)proteomics data. Optional in-source fragmentation and metabolomics experiments provide additional chemical information to guide the exploration of biosynthetic routes or phylogenetic relations. **b)** The first step— establishing the proteome-specific sugar components—performs mass binning of the complete set of fragmentation spectra acquired at very high mass resolution. Binned spectra thereby show highly comparable pattern across proteomes/species because those are the products of the universal set of amino acids (mirror bar chart). Carbohydrate fragments, however, have a characteristic chemical composition and, therefore, (usually) lay outside the amino acid composition space. Signals are automatically matched against a constructed database, consisting of over 3300 theoretical carbohydrate compositions. The procedure is exemplified by the heptose fragment (m/z 193.071), observed in the *Ca.* Kuenenia stuttgartiensis enrichment (meta)proteome. The heptose signal (193.071Da) is unique to Kuenenia, but the neighbouring mass peaks (−0.01Da/+0.026Da) are also observed in the non-glycosylated comparator proteome (small mirror bar chart) and are, therefore, amino acid related. **c)** The second step—termed ‘parent (ion) offset binning’—establishes the actual oligosaccharide chains which are linked to the proteins of individual strains. This is achieved by binning the mass deltas obtained after subtracting the fragment masses (fm) from their parent (ion) peptide mass (pm). Spectra containing the same carbohydrate fragments will, thereby, show repeatedly mass deltas consistent with the mass of the oligosaccharide modification. **d)** The established oligosaccharide chains are then integrated into the metaproteomic database search (using e.g. a metagenomics constructed databases), to identify the target proteins and related strains.

Because prokaryotes often produce species-unique carbohydrate derivatives, this strategy is highly advantageous as opposed to relying only on oxonium ion markers known from eukaryotic glycoproteins (e.g., HexNAc, Hex or NeuAc).

The second step—termed ‘parent offset binning’—establishes the oligosaccharide chains as modifying individual proteins (Figure 1D). Thereby, the peak mass lists of fragmentation spectra that contain carbohydrate components—and that (likely) derive therefore from oligosaccharide modified peptides—are now subtracted from the parent peptide mass. This approach generates mass deltas starting with the parent peptide mass at zero. Thereby, fragmentation spectra from glycopeptides will (repeatedly) show numbers consistent with the mass of the peptide-linked oligosaccharide chain(s). This process takes advantage of the predominant fragmentation of the carbohydrate chain over the peptide backbone when performing fragmentation by collision-induced dissociation, such as by higher-energy collisional dissociation (HCD) specific to Orbitrap mass spectrometers.^[25]^

Finally, binning parent offset mass deltas—that originate from spectra containing the same carbohydrate fragments—will provide a histogram of the oligosaccharide chain mass(es) present. This procedure relies on the presence of the y_0_ - ion in the fragmentation spectrum (unmodified, intact peptide fragment peak). Due to the strong fragmentation of the carbohydrate chains, however, this peak is very commonly observed in glycopeptide (CID/HCD) fragmentation spectra.^[25a, 25d, 25e]^

Moreover, owing to the oligosaccharide backbone fragmentation, binned spectra contain additional oligosaccharide fragments, therefore providing sequence and in some cases even linkage-type (N/O) information ^[25a, 25e]^ (Figure 3B, SI-DOC-D). Although prokaryotic protein glycosylation shows a large species and strain variability, the structural heterogeneity within one proteome is, generally, comparatively low. Hence, the binning approach is particularly for prokaryotic glycoproteomes a very useful procedure. Additionally, establishing the sugar components or oligosaccharide profiles does not require a proteome sequence database. Therefore, the (Matlab) data processing pipeline requires only a few minutes data processing time, e.g. when applied to single-run QE Orbitrap shotgun proteomics data. Finally, the thereby-established oligosaccharide profiles provide a database for recently developed high-performance glycopeptide annotation tools, ^[19b, 19c]^ or can be simply integrated as variable modifications into (multi-round) search approaches to identify modified proteins and target strains. (Figure 1E). Ultimately, this provides the common statistical parameters, such as peptide scores and false discovery rates. Most importantly, however, the determined chemical and compositional information guides physicochemical interpretation of the oligosaccharide chains, and the exploration of biosynthetic routes or phylogenetic relations.

### AN UNEXPECTEDLY COMPLEX ARRAY OF SUGARS

To verify the developed (‘sugar-miner’) approach for establishing sugar components directly from shotgun proteomics data, we processed a set of well-characterized control samples (Figure 2A, SI-DOC-C3:6). In so doing, every control proteome sample showed sugar components reflecting the known oligosaccharide structures of the respective species. Moreover, the theoretically constructed ‘carbohydrate composition space’, used to match carbohydrate fragments in the mass binning data, only marginally overlapped with peptide related fragments (e.g., those found in the *E. coli* comparator proteome). One of the few observed coincidences, however, included the compositional overlap of peptide-related fragments with the sugar composition of bacillosamine (diNAcBac) (Figure 2B, SI-DOC-B3:5). Furthermore, we processed all prokaryotic control samples also through the developed ‘parent offset binning’ procedure—which provides the oligosaccharide chains—to confirm the generic nature of this approach (SI-DOC-D3).

**FIGURE 2.**
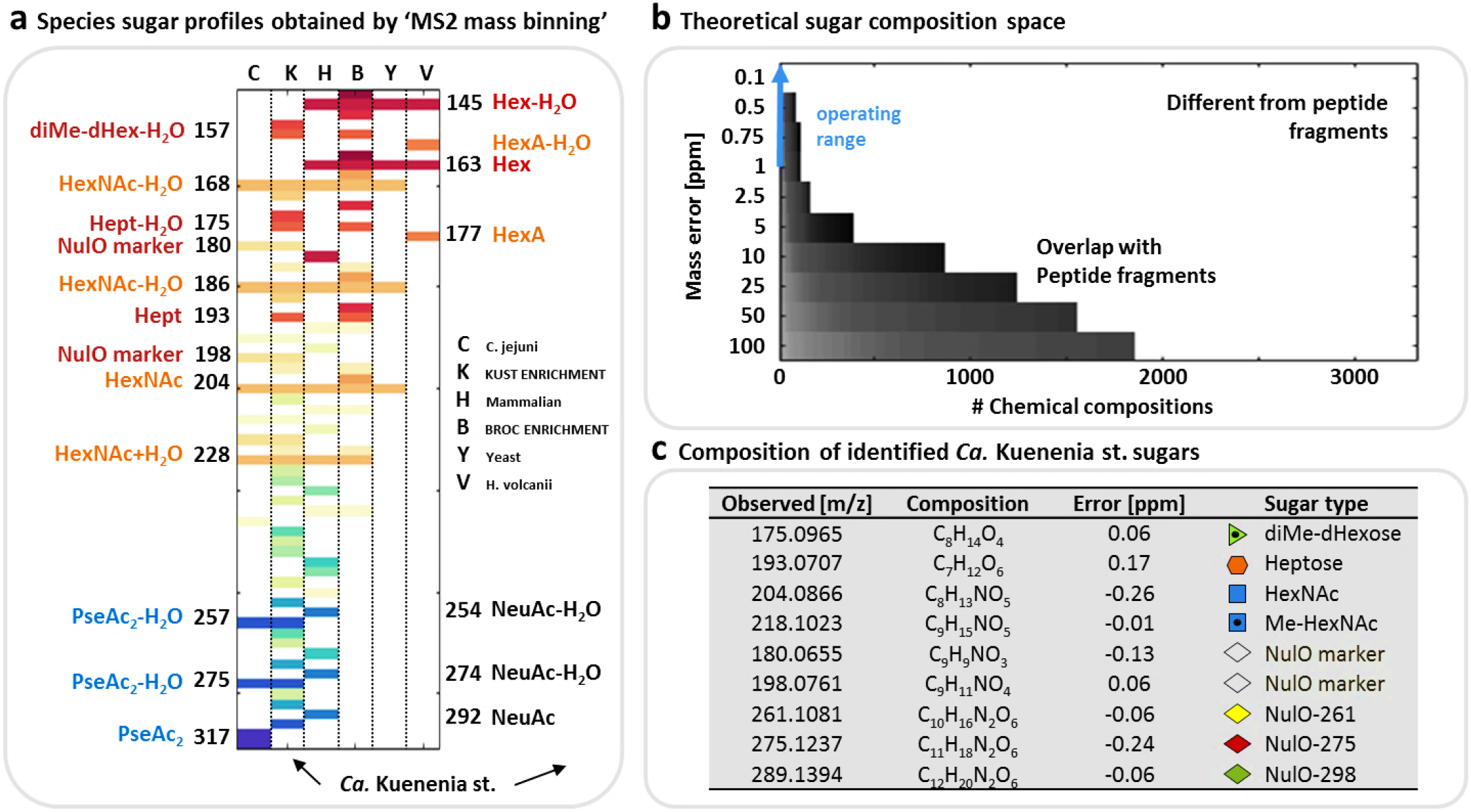
Carbohydrate profiles obtained from the anammox enrichment cultures (and reference samples) using the ‘MS2 mass binning’ approach. **a)** The graph (from left to right) shows the carbohydrate profiles (m/z values are distributed along the y-axis) from the *C. jejuni* sample (C, control)*, Ca.* Kuenenia stuttgartiensis enrichment culture (K), HEK protein sample (H, control), *Ca.* Brocadia sapporoensis enrichment culture (B), *S. cerevisiae* sample (Y, control) and *H. volcanii* sample (V, control). The carbohydrate profiles of the control samples established by the MS2 mass binning approach reflected the known oligosaccharide compositions of the individual species. Only sugar components which commonly provide only very low abundant or no carbohydrate related fragments (oxonium ions), such as deoxy hexoses, were not observed. NulO and HexNAc fragments are prominent in the *C. jejuni* proteomics sample, NeuAc/HexNAc and Hex related signals in the HEK derived sample, HexNAC- and hexose-related signals in the *S. cerevisiae* (yeast) proteomics sample, and hexuronic acid (HexA)- and hexose (Hex)-related signals in the *H. volcanii* sample. The latter was cultured at high salinity; therefore, the proteins appeared modified by only a single type of glycan structure. Fragments with the same colour indicate water loss clusters (−H_2_O), which are a characteristic consequence of the oligosaccharide chain fragmentation process, and therefore indicators for the discovery process. The mass compositions and sugar type annotations for the *Ca.* Kuenenia stuttgartiensis enrichment proteome are detailed in the table on the right. The abbreviations ‘NulO’ stand for nonulosonic acid, ‘NeuAc’ for N-Acetyl neuraminic acid; Pse’ for pseudaminic acid; ‘HexNAc’ for N-Acetyl-hexose amine; ‘Hept’ for heptose; ‘hex’ for hexose; ‘HexA’ for hexuronic acid; ‘dHex’ for deoxyhexose and ‘Me’ for methyl, respectively. **c) The constructed carbohydrate chemical composition space.** A large chemical composition space was constructed (>3300 theoretical compositions) used to assign chemical compositions to non-peptide related features. At high mass resolution (>100K) and accuracy (<1ppm), the mass overlap with amino acid-related fragments is considerably low. Mass recalibration using frequently observed amino acid fragments enables to operate at very high mass accuracy (blue arrow).

When exploring the sugar components found in the *Ca.* Kuenenia stuttgartiensis enrichment proteome, we observed a surprisingly diverse repertoire of carbohydrate fragments, including yet-undescribed nonulosonic acid derivatives as well as seven-carbon sugars, rarely observed in glycoproteins (Figure 2A:B, SI-DOC-C1). In addition, when investigating the related oligosaccharide profiles using the above-described ‘parent ion offset’ approach, we observed two completely unrelated types of oligosaccharide chains (Figure 3B, SI-DOC-D1). One type of oligosaccharides resembled the recently described N-acetyl-hexoseamine core, albeit containing nonulosonic acids not resolved in an earlier study (‘complex type structures’; X-type).^[14a]^ The second type of oligosaccharide structures consisted of homogeneous heptose chains (‘oligo-heptosidic’; O-type). More surprisingly, when integrating the oligosaccharide masses into a (multi-round) variable modification search using a metagenomics constructed database, both types of oligosaccharide structures were exclusively matched to the same surface layer protein (SLP) of *Ca.* Kuenenia stuttgartiensis (‘KUST_250_3’, SI-DOC-F1:3).

**FIGURE 3.**
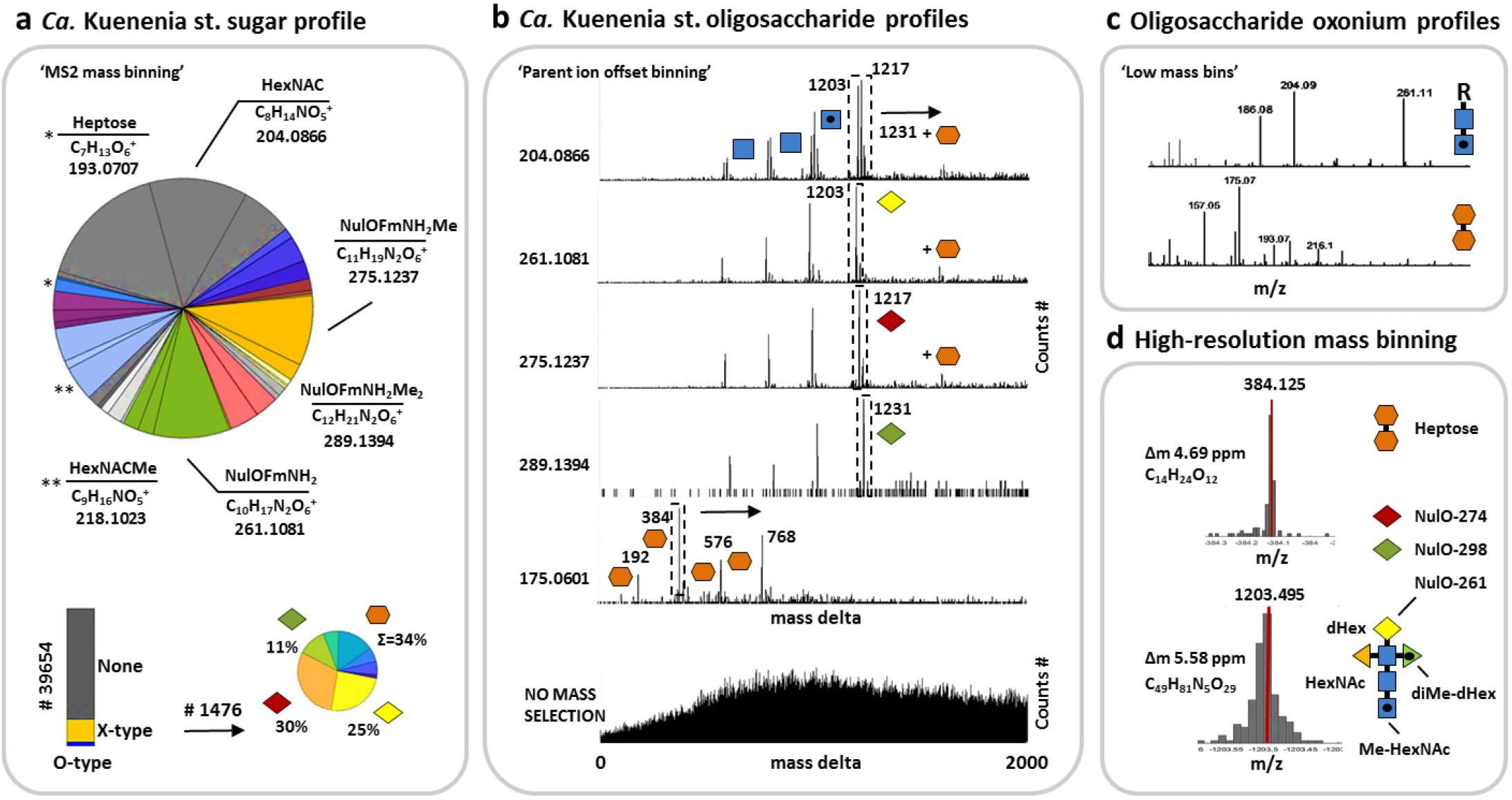
Outline of identified sugar components and oligosaccharide chain profiles for the *Ca.* Kuenenia stuttgartiensis enrichment culture. **a)** The large pie chart outlines the proportions of the carbohydrate fragments identified in the *Ca.* Kuenenia stuttgartiensis enrichment proteome using the MS2 mass binning approach. The lower charts depict the proportions of spectra containing carbohydrate related signals (oxonium ions) in the sequencing spectra, and the frequency of individual glycoforms across all spectra. **b)** The graphs outline the furthermore established oligosaccharide chains using ’parent ion offset binning’ of fragmentation spectra containing the identified carbohydrate fragments (graphs labelled with 204, 275, 289 and 175 m/z). Thereby, oligosaccharide chains appear in histograms as repeatedly occurring mass deltas. The same ‘parent ion offset’ approach applied to the complete, non-carbohydrate-filtered dataset, does not reveal any identifiable systematically reoccurring mass deltas (bottom graph). This revealed two completely unrelated oligosaccharide chains. The fragments 204, 261, 275 and 289 belong to variations of a complex type oligosaccharide with a HexNAc core structure (X-type). The fragment 175 (and 193) retrieved a second, fully unrelated heptose type oligosaccharide chain (O-type). Blue squares represent HexNAcs (methylations are depicted by a dot), yellow, red and green diamonds represent NulO variants, orange hexagons represent heptoses, orange triangles are deoxyhexoses, and green triangles (doted) are dimethyl-deoxyhexoses. Moreover, due to the predominant fragmentation of the oligosaccharide chains, nearly the complete sequence of the oligosaccharide can be derived. **c)** The histograms outline the (intensity normalized) low mass bins of fragmentation spectra where the complex type oligosaccharide (upper graph) or the oligo-heptosidic chains (O-type, lower graph) were identified. The thereby-observed sugar fragments correlate with the proposed composition of the individual carbohydrate chains (e.g. 204/261 for complex, or 175/193 for oligo-heptosidic). **d)** The histogram shows binning of mass deltas into very small bin sizes to establish the oligosaccharide compositions at very high mass accuracy (<7.5ppm).

Moreover, both oligosaccharides were apparently O-linked, and the complex type glycans followed the attachment motif GT/S, whereas no consensus glycosylation motif was identified for the ‘oligo-heptosidic’ chains (SI-DOC-G5:6). Interestingly, to the best of the author’s knowledge, a comparable complexity of unrelated glycosylation machineries targeting the same SLP simultaneously has only been observed before for archaea.^[6, 26]^ When comparing the results to a laboratory enrichment of *Ca.* Brocadia sapporoensis, another anammox species within the *Candidatus* Brocadiaceae family, a similar carbohydrate profile was observed, albeit lacking the nonulosonic acid-related fragments found in *Ca.* Kuenenia (Figure 2A, SI-DOC-C2). Furthermore, the observed sugar components established at least 4 different types of oligosaccharide chains (SI-DOC-D2). One was identical to a recently reported HexNAc core oligosaccharide (‘204’) ^[14b]^, another oligosaccharide structure contained characteristic hexose/heptose residues (‘163/193’), a third included characteristic methyl-deoxy hexose residues (‘161’), and the fourth was based on a yet-unidentified derivative (‘232’). When integrating the established oligosaccharide masses into the metaproteomic analysis using a specifically constructed metagenomic database, only the recently reported HexNAc core type oligosaccharide (‘204’) could be assigned to *Ca.* Brocadia sapporoensis, thereby exclusively modifying the putative SLP as also described in an earlier study.^[14b]^ The other types of oligosaccharide chains could be assigned to different proteins from Ignavibacteria bacterium OLB4 and Ignavibacteria bacterium UTCHB3, respectively, which both are commonly observed community members of anammox enrichment cultures^[27]^ (Figure 4A, SI-DOC-F4:6).

**FIGURE 4.**
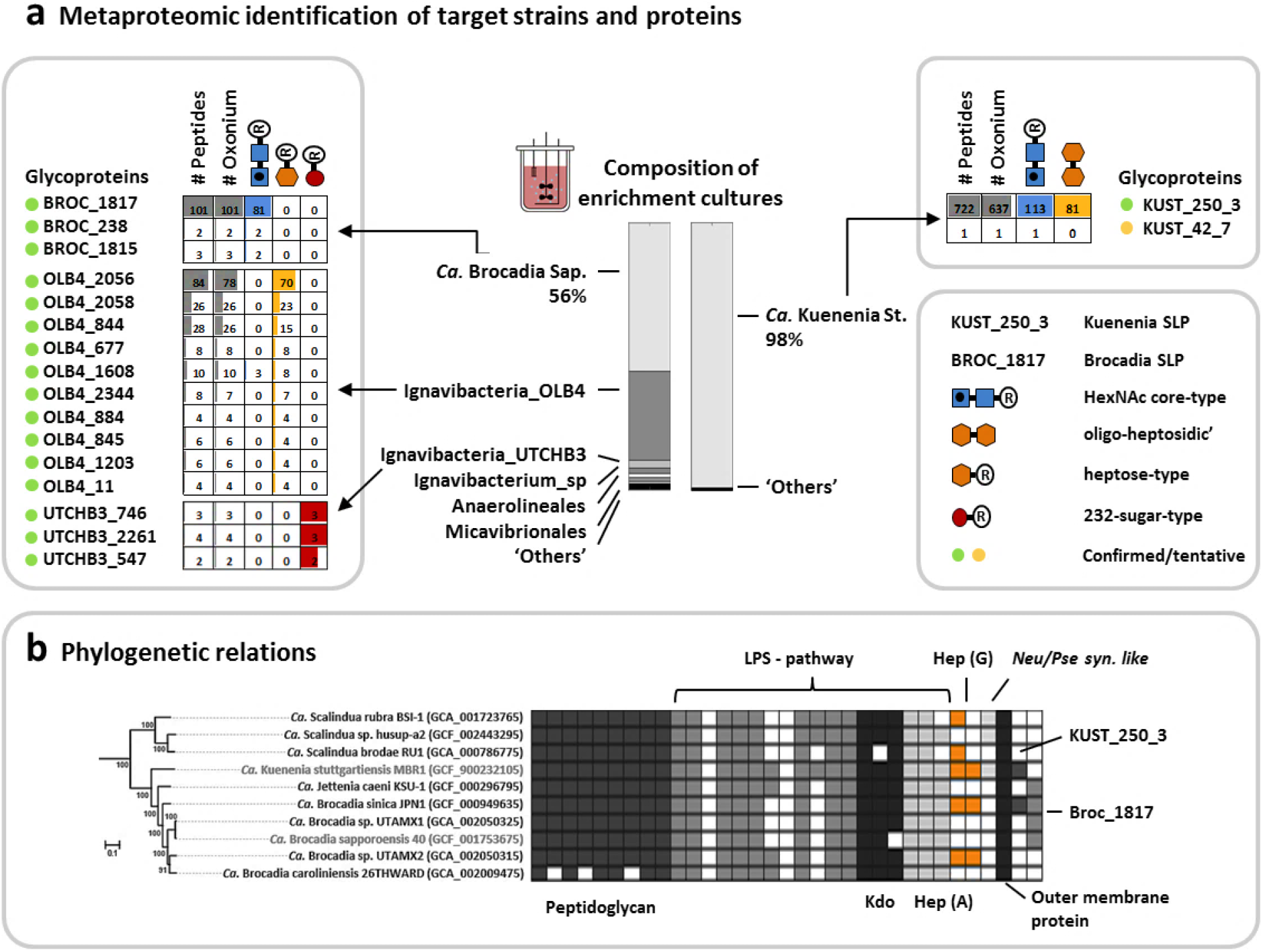
Glycosylated proteins and strains present in the explored anammox enrichment cultures. **a)** The metaproteomic analysis of the Kuenenia enrichment culture showed that approx. 98% of identified peptide sequences derived from *Ca.* Kuenenia stuttgartiensis (right bar). The vast majority of glycosylated peptides moreover could be assigned to the surface layer protein (KUST_250_3) of *Ca.* Kuenenia stuttgartiensis (right table). This confirmed that both types of oligosaccharides (‘complex-type’ and ‘oligo-heptosidic’) target the same surface layer protein simultaneously. Furthermore, the metaproteomic analysis of the Brocadia enrichment culture showed that approx. 56% of the peptide sequences derived from *Ca.* Brocadia sapporoensis, and significant other proportions to at least three different Ignavibacteria strains (left bar). A modification search, including the identified oligosaccharide chains, confirmed that the putative surface layer protein of *Ca.* Brocadia sapporoensis is modified by a HexNAc core type oligosaccharide (204, blue squares). A second type of oligosaccharide chain (161/193 = orange hexagons) was assigned to multiple proteins from Ignavibacteria bacterium OLB4, and a third type of oligosaccharide (232, red circles) to several proteins from Ignavibacteria bacterium UTCHB3 (left table). Only the top matches for every database search are shown in the graph. Proteins with peptide matches at all three levels (i–iii) were considered as confirmed (green circle; i=database search, ii=oxonium ions and iii=‘oligosaccharide mass deltas). ‘Peptides’ indicates the number of variable modification search matches (VM); ‘oxonium’ indicates the number of VM matches with additional oxonium ion identifications; ‘blue squares’ show the number of HexNAc core type oligosaccharide matches assigned to the same VM matches; ‘orange hexagon’ counts the number of heptose-type oligosaccharide matches that were also assigned to VM matches; ‘red circle’ counts the number of ‘232 sugar’-type oligosaccharide matches that were also assigned to the same VM matches. **b) The graph shows a maximum-likelihood phylogenetic tree, displaying the conservation of hypothesized genes required for producing the***Ca.* **Kuenenia stuttgartiensis surface layer**. The neuraminate synthase-like homologue found in the genome of *Ca.* Kuenenia stuttgartiensis (required to synthesize nonulosonic acids) appears more deeply rooted into the phylum of Planctomycetes, but was not present in the genome of *Ca.* Brocadia sapporoensis. Homologues to the GDP-heptose pathway, moreover, are exclusive to *Ca.* Kuenenia stuttgartiensis and *Ca.* Brocadia sinica. Most interestingly, the latter was also the only other strain possessing a homologue of the (highly acidic) surface layer protein found in *Ca.* Kuenenia stuttgartiensis. In contrast, the less acidic surface layer protein produced by *Ca.* Brocadia sapporoensis (and the ‘complex-type’ oligosaccharides), appear to be more common within the family of *Ca. Brocadiaceae*. Selected genes from peptidoglycan (PG) and lipopolysaccharide (LPS) pathways are widely present across the selected planctomycetal and *Ca. Brocadiaceae* genomes. The phylogenetic analysis was performed with the LG+F+I+G4 substitution model based on 37016 amino acid positions; the scale bar represents amino acid substitutions per site. Bootstrap values are listed as percentages at the branching points. The tree was rooted using *V. spinosum* DSM 4136 (GCF_000172155) as outgroup. Accession numbers for the genomes are indicated in brackets.

The established oligosaccharide chains for *Ca.* Kuenenia stuttgartiensis were also confirmed by orthogonal HILIC glycopeptide enrichment procedures combined with an open modification search (using Byonic), as proposed very recently^[22]^ (SI-DOC-D1). Moreover, we investigated an isolate of the *Ca.* Brocadia sapporoensis SLP separately, to confirm that it is modified by only one type of oligosaccharide (‘204’). To this end, we performed a conventional SDS-PAGE^[14b]^, followed by in-gel proteolytic digestion and mass spectrometric analysis. The shotgun proteomic data were then processed by the developed pipeline. This showed exactly the same HexNAc-type oligosaccharide (‘204’) as identified from the large-scale data and confirmed the lack of ‘oligo-heptosidic’ chains found in *Ca.* Kuenenia stuttgartiensis (SI-DOC-D2). In summary, both strains—*Ca.* Kuenenia stuttgartiensis and *Ca.* Brocadia sapporoensis—share the HexNAc core type oligosaccharides (X-type), but they differ in regard to the presence of terminal nonulosonic acids and the second ‘oligo-heptosidic’ chains. Nonetheless, the physiological importance of this extensive and complex surface glycosylation discovered for *Ca.* Kuenenia stuttgartiensis remained unknown to this point.

### A FINE BALANCE BETWEEN CELLULAR PROTECTION AND SUBSTRATE AQUISITION

While *Ca.* Kuenenia stuttgartiensis gains energy by oxidizing positively charged ammonium (NH_4_^+^) with negatively charged nitrite (NO_2_^−^) to yield dinitrogen gas (N_2_)^[11a]^, it features high metabolic versatility^[28]^. Yet, little is known about how anammox bacteria adapt to fluctuating substrates availabilities. Only recently, Smeulders et al. started to shed light on this aspect and revealed the capability of anammox bacteria to differentially express their multiple transporter genes in response to substrate limitations.^[29]^ However, whether anammox bacteria undertake additional adaptations, e.g. physiological, in response to its metabolic requirements, has not been shown to date.

Almost all archaea and many bacteria are entirely covered by surface layer proteins (SLPs).^[30]^ Moreover, SLPs frequently show either acidic or basic isoelectric points due to their propensity for charged amino acids.^[31]^ Li et al.^[31]^ demonstrated that the negatively charged SLPs of ammonia-oxidizing archaea (AOA) support the acquisition of ammonium (NH_4_ ^+^), which supposedly enables AOA to be competitive in nutrient-limited ecosystems.^[31]^ Moreover, Li et al. showed that the opposite case of a positively charged surface layer would create a nutrient barrier, preventing NH_4_ ^+^ from passing through the surface pores into the pseudo-periplasmic space.^[31–32]^

*Ca.* Kuenenia stuttgartiensis possesses a very comparable (hexagonally arranged) SLP array covering the entire bacterial cell^[14c]^, and the specific SLP has a particularly acidic (predicted) isoelectric point of approx. 4.25 and a net charge of −60 at physiological pH (Figure 5A, SI-DOC-G1:2). Intriguingly, however, anammox bacteria not only depend on the acquisition of ammonium but also require negatively charged nitrite. These findings raise the question of why *Ca.* Kuenenia stuttgartiensis possesses such a highly acidic SLP, and whether *Ca.* Kuenenia stuttgartiensis (or anammox bacteria in general) evolved additional modulations to avoid interference with substrate acquisition. This is of particular importance as nitrite is commonly the limiting substrate in engineered and natural ecosystems.

**FIGURE 5.**
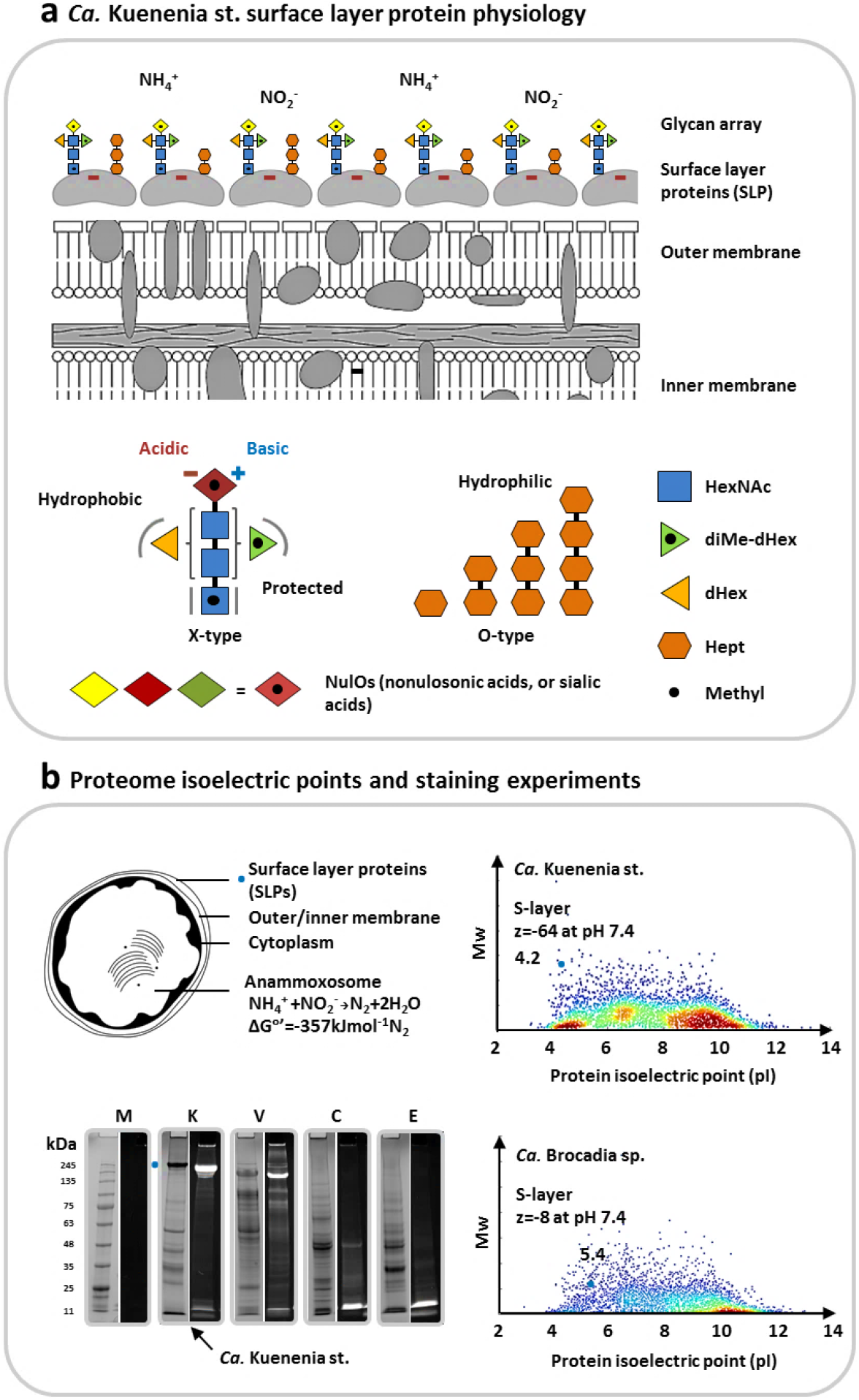
Physiology of the *Ca.* Kuenenia stuttgartiensis surface layer protein (SLP) and oligosaccharides. **a)** The surface layer protein (SLP) of *Ca.* Kuenenia stuttgartiensis is densely covered by two entirely different types of oligosaccharides (‘X-type’ and ‘O-type’). The dense layer supposedly provides shielding of the very acidic SLP. Because anammox bacteria gain their metabolic energy through ammonia oxidation—which requires the acquisition of positively charged NH4+ and negatively charged NO_2_^−^-—a ‘charge-balanced’ surface appears highly advantageous. Interestingly, the investigated *Ca.* Kuenenia stuttgartiensis strain produces nonulosonic acids (NulOs) which possess an unmasked amine. Those have the potential to counterbalance the carboxylic acid groups. **b)** The *Ca.* Kuenenia stuttgartiensis surface layer protein shows a predicted pI (isoelectric point) of approx. 4.25 and a net charge of approx. −60 at physiological pH. In fact, the surface layer protein is one of the most acidic proteins of the complete *Ca.* Kuenenia stuttgartiensis proteome. On the other hand, the putative surface layer protein of *Ca.* Brocadia sapporoensis has a predicted pI of only 5.4 and a substantially lower net charge of approx. −8 at physiological pH. Moreover, *Ca.* Brocadia, uses also only a related form of the complex-type oligosaccharide to cover its much less acidic surface layer protein. The SDS-PAGE analyses show protein and sugar staining for the protein extracts from *Ca.* Kuenenia stuttgartiensis and the additional control strains *H. volcanii* (glycan-positive control), *C. jejuni* (glycan-positive control) and *E. coli K12* (glycan-negative control). The left lanes each show the total protein staining (‘P’; Brilliant Blue G staining solution), whereas the right lanes each show the carbohydrate staining (‘C’; Pro-Q 488 Emerald staining kit).

Interestingly, Hu et al. recently revealed the ability of anammox to grow on neutral nitric oxide (NO) instead of nitrite. As NO was likely the first oxidized nitrogen form present on early Earth^[33]^, it is tempting to hypothesize that the acquisition of the highly acidic SLPs—analogue to AOA, which only depend on positively charged ammonium—preceded the ability of anammox bacteria to utilise negatively charged nitrite. Nevertheless, this could not explain how current anammox acquire negatively charged nitrite. Finally, the development of the here-discovered dense layer of (charge-balanced) oligosaccharides, may have provided sufficient shielding of the highly acidic protein layer to enable the acquisition of negatively charged nitrite without potential interference (Figure 5B).

However, the investigated *Ca.* Kuenenia stuttgartiensis strain produces complex oligosaccharides, including nonulosonic acids. Those sugars are commonly associated with enhancing cellular protection through surface diversification (e.g., in response to bacteriophage recognition).^[8a, 34]^ Yet, nonulosonic acids are highly acidic sugars and support a negative surface charge.^[8a, 35]^ Thus, although those sugars may be of advantage for cellular protection, it seems counterintuitive to invest cellular energy into an (even more) negatively charged surface layer.

Nevertheless, carboxylic acids can be chemically modified or balanced through basic counterparts^[36],[35]^; for example, through esterification of the carboxylic acid groups or by free amines, which are otherwise present in alkylated forms.^[36–37]^ Surprisingly, the chemical composition of the nonulosonic acids, additional in-source fragmentation and labelling experiments indeed indicate the presence of an unmasked amine (Figure 5C, SI-DOC-E1:2,8). Those unequivocally have the potential to counterbalance the neighbouring, highly acidic carboxylic acids.^[35]^ While zwitterionic sugar modifications have been detected (e.g., recently in glycans of non-vertebrates)^[38]^, nonulosonic acids with free amines have been only rarely observed but were described, for example, in nerve and cancer cells under certain conditions.^[35, 37]^

In summary, *Ca.* Kuenenia stuttgartiensis employs two glycosylation machineries to cover its SLP with a dense array of oligosaccharides. The charge-balanced oligosaccharides have the potential to shield the particularly acidic surface layer protein. Furthermore, the investigated *Ca.* Kuenenia stuttgartiensis strain produces complex oligosaccharides, including ‘unusual’ nonulosonic acids, which further underlines the apparent necessity to ‘charge-balance’ the entire bacterial cell surface layer. Alternatively, the evolutionary loss of the SLP would have likewise resolved any potential substrate acquisition conflicts. However, this would have also significantly reduced cellular protection or other important features of SLPs required to survive in a competitive environment.^[30]^

To place our findings in the context of the broader anammox physiology, we also investigated the *Ca.* Brocadia sapporoensis surface layer protein (SLP). This revealed a substantially smaller and less acidic surface layer protein (predicted pI_~_5.4), which furthermore is modified by only one type of oligosaccharide (Figure 5A, SI-DOC-G3:4). These differences are likely to contribute to the reported divergences between strains of the genus *Ca.* Kuenenia and *Ca.* Brocadia, for example in regard to substrate affinity (i.e., the ability to thrive at low concentrations) or the tendency to grow in free-living planktonic form. ^[39]^

### PHYLOGENETIC CELL SURFACE LAYER DIVERSITY

Additional metabolomic analysis of the *Ca.* Kuenenia stuttgartiensis enrichment culture indicated that the nonulosonic acids are cytosine monophosphate (CMP) activated (the common activation route for nonulosonic acids), and that heptoses are present in adenosine diphosphate (ADP) as well as guanosine diphosphate (GDP) activated forms. The ADP-activated form is the common substrate used to synthesize the conserved lipopolysaccharide (LPS) core, the corresponding biosynthetic genes of which are present in *Ca.* Kuenenia stuttgartiensis.^[40]^ Interestingly, on the other hand, GDP-activated heptoses have been described as substrates for the SLP glycosylation of the Gram-positive *Bacillus thermoaerophilus* (SI-DOC-E7).^[41]^ To explore potential conserved elements, we investigated the hypothesized genes required for producing the *Ca.* Kuenenia stuttgartiensis surface layer across selected planctomycetal genomes (Figure 4B, SI-DOC-J). This showed that a putative neuraminate synthase-like (or Pse/Leg synthase) found in *Ca.* Kuenenia stuttgartiensis genome, required for biosynthesis nonulosonic acids, is more deeply rooted into the phylum Planctomycetes. However, homologues to the GDP-heptose pathway, in particular the D-glycero-alpha-D-manno-heptose 1-phosphate guanylyl transferase, seem nearly exclusive *to Ca.* Kuenenia stuttgartiensis. Most interestingly, the only other homologue was found in *Ca.* Brocadia sinica, which also possesses a homologue of the SLP found in *Ca.* Kuenenia stuttgartiensis (NCBI CAJ72259). This finding suggests that the (more) acidic SLP and the O-type oligosaccharides may have been acquired simultaneously. In contrast, the HexNAc core type oligosaccharide (X-type), a related form of which has also been experimentally confirmed in *Ca.* Brocadia sapporoensis^[14b]^, and the less acidic SLP (Broc_1817) seem to represent a more widely distributed feature within the family *Ca.* Brocadiaceae.

However, we cannot exclude the possibility that the observed O-type glycans are synthesized from ADP-heptose substrates. Required heptosyltransferases, which are apparently present in associated community members such as the Ignavibacteria species that produce related glycans (Figure 4A) but lack the genes of the GDP-heptose pathway, could be acquired through horizontal gene transfer, which is a rather common process for carbohydrate-active enzymes.^[42]^

## CONCLUSIONS

Protein glycosylation – one of the most abundant protein modifications – is fundamental to all three domains of life. However, the enormous chemical diversity and the large strain variability in prokaryotes makes their exploration exceptionally challenging. Therefore, the current understanding of their biological significance remains largely unknown.

Here, we provide a universal procedure to explore prokaryotic protein glycosylation from non-pure cultures. The approach provides insights into the chemical identity of novel sugar components and establishes the related protein linked oligosaccharide chains from large-scale (meta)proteomics data directly.

We applied the approach to an anammox bacteria enrichment culture, and resolved a remarkably complex array of surface layer oligosaccharides. The identified glycans are produced by two entirely unrelated glycosylation machineries and densely cover a very acidic surface layer protein. Moreover, the investigated anammox strain accomplished modulation of highly specialized sugars, presumably in response to its metabolic requirements—the anaerobic oxidation of ammonium—which depends on the acquisition of substrates of opposite charge.

Ultimately, the physicochemical interpretation of the discovered spectrum of oligosaccharides suggests a broader link to the evolution of charged surface layer proteins and the ‘metabolic lifestyle’ in anammox bacteria.

## Supporting information

Supplemental FILE INDEX

## SUPPLEMENTAL INFORMATION

Electronic supplementary information material is available: additional supporting method validation procedures including figures and tables, and supporting raw data tables. The Matlab ‘Sugar-miner’ and ‘Glyco-mod-pro’ scripts are freely available upon request.

## ACKNOWLEDGEMENTS

We acknowledge Laura van Niftrik for reading the manuscript and providing constructive feedback. We further would like to acknowledge Claire Chassagne for discussions on surface charges, Guylaine Nuijten and Katinka van de Pas-Schoonen for anammox biomass sampling, technical assistance and reactor care, and Ben Abbas for the support with DNA extraction. The authors acknowledge the SIAM consortium and the TU Delft startup fund for funding. Additionally, SL was supported by a NWO VIDI grant (016.Vidi.189.050), and ML was supported by a Marie Skłodowska-Curie Individual Fellowship (752992), and a VENI grant from the Dutch Research Council (NWO, VI.Veni.192.252).

## MATERIALS AND METHODS

### (A) SAMPLE SOURCES MICROBES

#### Anammox lab-scale enrichment cultures

*Ca.* Kuenenia stuttgartiensis was enriched as planktonic cells (approx. 90% relative abundance based on metagenomic data) at 30°C in a continuous-flow bioreactor equipped with a custom-made microfiltration module (pore size of 0.1μm) as described elsewhere.^[43]^ The reactor was fed with mineral medium^[43]^ supplemented with 45mM of ammonium and nitrite. Nitrite concentrations in the effluent were always below detection limit (nitrite test strips MQuant, Merck, Darmstadt, Germany). Anoxic conditions were maintained via continuous sparging with Ar/CO2 (95%/5% v/v) at a rate of 10ml/min. The reactor hydraulic and solids retention times were approximately 1.9 and 10.5 days, respectively, and the resulting steady-state OD600 was 1.0-1.1. The pH was controlled at 7.3 with a 1M KHCO_3_ solution. The flocculent *Ca.* Brocadia sapporoensis enrichment was maintained at 30°C under anoxic conditions using an identical continuous flow bioreactor with a custom-made microfiltration module (pore size of 0.1μm) fed with a concentrated media of 60mM ammonium and nitrite as originally described by Lotti et al., 2014.^[44]^

#### Comparators strain cultures

E coli K12, *H. volcanii* (cultured at high salinity, >3.5M NaCl), *C. jejuni and S. cerevisiae* control samples were cultured and harvested as described by Kleikamp et al., 2020.^[8a]^

### (B) MASS SPECTROMETRY BASED PROTEOMICS

#### Cell lysis and protein extraction

A modified protocol from Kleikamp et al. was used to prepare whole protein extracts.^[45]^ Briefly, 25mg biomass (wet weight) were collected in an Eppendorf tube and solubilized in a suspension solution consisting of 200µL B-PER reagent (78243, Thermo Scientific) and 200µL TEAB buffer (50mM TEAB, 1% (w/w) NaDOC, adjusted to pH 8.0) including 0.2μL protease inhibitor (P8215, Sigma Aldrich). Furthermore, 0.1g of glass beads (acid, washed, approx. 100μm diameter, G4649-10G, Sigma Aldrich) were added and cells were disrupted using 3 cycles of bead beating on a vortex for 30 seconds followed by cooling on ice for 30 seconds in-between cycles. In the following, a freeze/thaw step was performed by freezing the suspension at −80°C for 15 minutes and thawing under shaking at elevated temperature using an Eppendorf incubator (ThermoMixer™). The cell debris was pelleted by centrifugation using a bench top centrifuge at max speed, under cooling for 10 minutes. The supernatant was transferred to a new Eppendorf tube and kept at 4°C until further processed. Protein was precipitated by adding 1 volume of TCA (trichloroacetic acid) to 4 volumes of supernatant. The solution was incubated at 4°C for 10 minutes and subsequently pelleted at 14.000rmp for 10 minutes. The obtained protein precipitate was washed twice using 250μL ice cold acetone. The protein pellet was dissolved in 100μL of 200mM ammonium bicarbonate containing 6M Urea to a final concentration of approximately 100μg/μL. To 100μL protein solution, 30μL of a 10mM DTT solution were added and incubated at 37°C for 1 hour. In the following, 30μL of a freshly prepared 20mM IAA solution was added and incubated in the dark for 30 minutes. The solution was diluted to below 1M Urea using 200mM bicarbonate buffer and an aliquot of approximately 25μg protein were digested using sequencing grade Trypsin (V511A, Promega) at 37°C over-night (Trypsin to protein ratio of approx. 1:50). Finally, protein digests were then further desalted using an Oasis HLB 96 well plate (WAT058951, Waters) according to the manufacturer protocols. The purified peptide eluate was dried using a speed-vac concentrator.

#### ZIC-HILIC solid phase extraction of glycopeptides

SPEs (ZIC-tip, P/N 2942-660, 200μL, 10mg, 50μm, di2chrom, Germany) were washed 2x with 200μL H_2_O and further equilibrated with 2×200μL 80% AcN plus 0.1% TFA. To 7μL of sample 63μL of 85% AcN, 0.1% TFA was added and the sample was loaded on the tips (in 3 cycles) and washed by 70μL, 80% AcN, 0.1% TFA. The retained glycopeptides were eluted using 100μL HPLC grade H_2_O, and finally were speed-vac dried for further analysis.

#### SDS-PAGE, glycostaining and in-gel proteolytic digestion

Whole cell lysate protein extracts from *Ca.* Kuenenia stuttgartiensis enrichment, *Ca.* Brocadia sapporoensis enrichment, *C. jejuni*, *H. volcanii*, *E. coli K12* whole cell lysates were analysed using 10% Mini-PROTEAN Protein Gels (4568034, Bio-Rad, Laemmli buffer S3401-VL, Sigma Aldrich). One part was stained using Coomassie Blue for total protein content (Brilliant Blue G solution, Sigma Aldrich) and a second part was stained and analysed for glycoproteins using the Pro-Q 488 Emerald Glycoprotein staining kit (P33375, Thermo/Invitrogen), according to the manufacturers protocol. The enrichment, SDS-PAGE analysis and in-gel digestion of the putative *Ca.* Brocadia sapporoensis surface layer protein was performed as described in Boleij et al., 2018.^[14b]^

#### Whole cell lysate shotgun proteomics and related experiments

The vacuum dried peptide fractions were resuspended in H_2_O containing 3% acetonitrile and 0.1% formic acid under careful vortexing. An aliquot corresponding to approx. 250ng protein digest was each analysed using an one dimensional shotgun proteomics approach.^[46]^ Briefly, samples were injected to a nano-liquid-chromatography system consisting of an ESAY nano LC 1200, equipped with an Acclaim PepMap RSLC RPC18 separation column (50μmx150mm, 2μm and 100Å), and an QE plus Orbitrap mass spectrometer (Thermo Scientific, Germany). Unless otherwise specified, the flow rate was maintained at 300nL/min over a linear gradient using H_2_O containing 0.1% formic acid as solvent A, and 80% acetonitrile in H_2_O and 0.1% formic acid as solvent B. Solvent gradients and acquisition modes used for the individual experiments are detailed in the following. A) High-resolution MS2 mass binning experiments: The peptides were analysed using a gradient from 4% to 30% solvent B over 32.5 minutes, and finally to 70% solvent B over 12.5 minutes. The Orbitrap was operated in data-dependent acquisition (DDA) mode acquiring peptide signals form 500–1500 m/z at 70K resolution with an AGC target of 3e6. The top 10 signals were isolated with a 1.6m/z window and fragmented using a NCE of 30. The AGC target was set to 2e5, at a max IT of 100ms, a fixed first mass of 120, and a resolution of 140K. Dynamic exclusion was set to 20 seconds. Mass peaks with unassigned charge state, singly, 7 and >7, were excluded from fragmentation. B) Whole cell lysate shotgun glycoproteomics: The peptides were analysed using a gradient from 5% to 30% solvent B over 85 minutes, and finally to 75% B over 25minutes. The Orbitrap was operated in data-dependent acquisition (DDA) mode acquiring peptide signals form 550–1500 m/z at 70K resolution with an AGC target of 3e6. The top 10 signals were isolated with a 2.0m/z window and fragmented using a NCE of 28. The AGC target was set to 2e5, at a max IT of 75/54ms, a fixed first mass of 120, an isolation offset of 0.1m/z, and a resolution of 17K. Dynamic exclusion was set to 20 seconds. Mass peaks with unassigned charge state, singly, 6 and >6, were excluded from fragmentation. C) Whole cell lysate shotgun proteomics: The peptides were analysed using a gradient from 5% to 30% solvent B over 85 minutes, and finally to 75% B over 25 minutes. The Orbitrap was operated in data-dependent acquisition (DDA) mode acquiring peptide signals form 385– 1250m/z at 70K resolution with an AGC target of 3e6. The top 10 signals were isolated with a 2.0m/z window and fragmented using a NCE of 28. The AGC target was set to 2e5, at a max IT of 75/54ms, a fixed first mass of 120, an isolation offset of 0.1m/z, and a resolution of 17K. Dynamic exclusion was set to 60 seconds. Mass peaks with unassigned charge state, singly, 6 and >6, were excluded from fragmentation. D) Analysis of in-gel digested proteins: The peptides were analysed using a gradient from 5% to 25% solvent B over 25 minutes, and finally to 60% solvent B over 10 minutes. The flow rate was maintained at 350nL/min. The Orbitrap was operated in data-dependent acquisition (DDA) mode acquiring peptide signals form 400– 1400m/z at 70K resolution with an AGC target of 3e6. The top 10 signals were isolated with a 1.6m/z window and fragmented using a NCE of 28. The AGC target was set to 5e4, at a max IT of 150ms, a fixed first mass of 120, an isolation offset of 0.1m/z, and a resolution of 17K. Dynamic exclusion was set to 60 seconds. Mass peaks with an unassigned charge state, singly, 6 and >6, were excluded from fragmentation. E) In-source fragmentation experiments: The peptides were analysed using a gradient from 6% to 30% solvent B over 40 minutes, and finally to 60% B over 15 minutes. The flow rate was maintained at 350nL/min. The Orbitrap was operated in positive ionisation mode acquiring signals alternating between PRM and Full MS-SIM mode. The PRM mode was performed at an in-source CID of 75eV, isolating the target sugar fragments with an isolation window of 0.4m/z, at 0.1m/z isolation offset and a loop count of 9. Fragmentation was performed using a NCE of 25, acquiring fragments at a resolution of 70K, using an AGC target of 5e5 and a max IT of 150ms. The fixed lowest mass was set to 50m/z. The Full MS – SIM mode was operated with an in-source CID of 75eV, acquiring full scan mass spectra at 70K resolution, at an AGC target of 3e6 and an max IT of 60ms, over a mass range of 140– 1400m/z. PRM carbohydrate fragment targets for the *Ca.* Kuenenia stuttgartiensis enrichment were set to 193, 175, 147, 204, 218, 261, 275, 163, 407; for the *Ca.* Brocadia sapporoensis enrichment to 147, 204, 218, 232, 334, 133, 352; and for the mammalian protein control sample to 147, 204, 292, 163.

#### PEAKS database search

Whole cell lysate shotgun proteomics raw data (see B/C) were analysed using PEAKS Studio X (Bioinformatics Solutions Inc., Canada) against a metagenomics constructed database. Database search was performed employing a two-round search strategy, where the first round was used to construct a focused protein sequence database, thereby allowing peptide spectrum matches up to 5% false discovery rate and protein matches without unique peptide assignments. The database search was performed allowing 50ppm mass error, 0.01Da fragment error tolerance, considering 2 missed cleavages, oxidation/deamination as variable modifications and carbamidomethylation as fixed modification. The cRAP protein sequences were downloaded from ftp://ftp.thegpm.org/fasta/cRAP. The second round search was performed including the identified oligosaccharides masses as variable modifications, allowing up to 2 variable modifications per peptide. Peptide spectra were filtered against 0.1% false discovery rate and reported protein identifications required ≥1 unique peptides, where protein identifications with ≥2 unique peptides were considered as significant.

#### Verification of glycopeptide spectrum matches

Matlab R2017b was further used to score glycopeptide spectrum matches for the additional presence of expected oxonium ions or for whether glycan structures have been identified in the same scans by ‘parent ion offset’ binning. Only proteins/species which provided spectra showing variable modification search identifications, oxonium ion and structural identifications in the same spectra, were considered as confirmed matches.

#### BYONIC open modification search

Confirmatory open modification search was performed on HILIC enriched glycopeptides using Byonic (Protein Metrics Inc). Open modification search was performed on all peptides using the wildcard search option, considering a modification mass range of 0–2000Da, and the target amino acids N/S/T/Y. The search included carbamido-methylation as fixed modification, allowed up to 2 missed cleavages, a precursor mass and a fragment mass error tolerance of 20ppm. The maximum precursor mass was set to 8000, where the precursor and charge assignments were computed form the MS1 spectrum. The maximum number of precursors was set to 1 and the smoothing width was set to 0.005m/z. The search was performed using the focused sequence database generated by the PEAKS first round search described above, and by including decoys and an automatic score cut off. The protein identifications were filtered for 1% false discovery rates, except stated otherwise. The peptide spectrum match list was imported into Matlab R2017b, where mass shifts from peptides spectrum matches with a Log Prob <1 were binned at 0.1Da mass windows using the ‘histogram’ function to create a frequency histogram. Mass shift peaks of the histogram were annotated using the ‘mspeaks’ function, following intensity normalisation and detection of bin counts.

#### Data availability

The mass spectrometry proteomics raw data have been deposited in the ProteomeXchange consortium database with the dataset identifier PXD021600.

### (C) GLYCOSYLATION ANALYSIS

#### ‘Sugar-miner’

Identification of strain specific carbohydrate fragments was established by the following steps: (A) mass binning of very high-resolution shotgun proteomics data; (B) establishing a theoretical chemical composition space for carbohydrate fragments; (C) annotation of (non-peptide) mass peaks with possible carbohydrate compositions; (D) verifying the annotations by investigating the presence of ‘water-loss clusters’, and by determining the co-occurrence (correlation) of the cluster peaks across the entire proteomics run. More specifically: A) Mass spectrometric shotgun raw data acquired at very high resolution (140K) were converted using peak picking ‘vendor’ into ‘.mgf’ files considering only second-level scans using the msConvertGUI tool (ProteoWizard). For the comparator *E coli* K12 sample an additional absolute int. threshold peak filter of 25K was applied. The ‘mgf’ files were imported into the Matlab environment using the Matlab ‘textscan’ function. The mass peaks from all second-level (MS2) scans were further combined into a single matrix, and masses in the range from 110–325m/z were binned into 0.0001m/z windows using the ‘histcounts’ function. The obtained raw traces were further corrected for mass drifts by alignment to known amino acid fragment peaks (147.1128; 175.1190; 201.1234; 215.1390; 228.1343; 258.1448; 292.1292) using the ‘msalign’ function, and further normalised to ‘100’ using the ‘msnorm’ function. The thereby generated ‘raw traces’ were converted into (centroided) peak lists using the ‘mspeaks’ function, employing a ‘heightfilter’ of 0.02% (relative to the largest peak). The same procedure (except using a relative ‘heightfilter’ of 0.1%) was applied simultaneously to the *E. coli* K12 non-glycosylated comparator strain dataset. B) An empirical carbohydrate fragment composition space was constructed considering the elemental composition space C_5-14_H_4-28_N_0-2_O_2-12_S_0-1_. The compositions were further filtered for realistic structures evaluating C/H and CO/N ratios, the degree of unsaturation (DBEs), the mass defect and the min/max absolute masses. For details see SI-DOC-B2. C) To identify possible carbohydrate signals in the sample data, the established sample peak list was matched with the empirical composition space at a mass tolerance of 0.75ppm. Mass peaks also present in the non-glycosylated *E. coli* K12 comparator at a relative level >0.5 were considered as amino-acid-related. D) The established carbohydrate fragment candidates were further evaluated for the presence of ‘water-loss clusters’ (−18.01 mass deltas between fragments), commonly observed when fragmenting carbohydrate compounds. To ensure that the assigned ‘water-loss clusters/pairs’ are part of the same parent structure, the clusters were evaluated for co-occurrence within the same scans across the shotgun proteomics dataset. For this, the ‘conventional mass resolution’ shotgun dataset (of the same sample) was converted to ‘mzXML’ using the msConvertGUI tool (ProteoWizard), considering first-and second-level scans. The ‘mzXML’ file was imported into the Matlab environment using the ‘mzxmlread’ function. The obtained ‘mzxmlstruct’ structure was processed using ‘mzxml2peaks’ and ‘arrayfun’ to extract first- and second-level spectral information. By doing so, an extracted ion chromatogram was collected (within +/-7.5ppm mass window) for every carbohydrate candidate, containing information about the occurrence across the scans. To evaluate the degree of co-occurrence (=correlation) between individual carbohydrate fragments, the matrix was further converted into a ‘correlation matrix’, by dividing the total occurrence by the number of scans shared between fragments. A correlation of ‘1’ indicates that all scans are shared, where a correlation of ‘0’ means that two carbohydrate fragments don’t share any scans. To ovoid accumulation of background signals, a minimum intensity of 5E4 was required. Only ‘water-loss pairs/clusters’ with a correlation >0 between, and >0.5 within the entire ‘water-loss cluster’ were considered for further processing. To avoid accumulation of background signals, the carbohydrate signals required a minimum relative intensity (0.01) or a minimum number of counts (10) across the entire dataset. The established carbohydrate compounds were exported to a ‘xlsx’ table.

#### ‘Glyco-mod-pro’

Establishing the oligosaccharide profiles as modifying the proteins is performed by the following steps:

(A) identifying scans in a shotgun proteomics run which contain the identified carbohydrate fragments; (B) calculating the ‘parent ion offsets numbers’ and (C) binning the ‘offset numbers’ to identify the reoccurring mass deltas (=oligosaccharide chains). More specifically: A) Mass spectrometric raw files of ‘conventional’ shotgun proteomics analysis runs were converted into ‘.mzXML’ files using the msConvertGUI tool (ProteoWizard). Files were further imported into the Matlab environment using the ‘mzxmlread’ function. Furthermore, a table of the exact masses of the identified carbohydrate fragments as established by the previous ‘sugar-mining’ step, was provided as ‘.xlsx’ table and further imported into the Matlab environment using the ‘xlsread‘ function. The constructed ‘mzxmlstruct’ structure was further processed using the ‘mzxml2peaks’ and ‘arrayfun’ functions to extract first- and second-level spectral information. A matrix was created containing mass/charge (m/z) values, ion intensities, scan numbers (and related parameters) of mass peaks matching the identified carbohydrate fragments, within a tolerance of +/− 7.5ppm. B) The ‘parent ion offset’ was further calculated for every second-level (MS2) spectrum containing a particular carbohydrate fragment. Briefly, for a particular fragment, all scans containing the carbohydrate fragment were collected. Scans with an unique parent ion mass (2 digits) were processed one by one, using the Matlab ‘for’ loop functions. First, the complete second-level (MS2) spectrum was extracted. Peaks with a mass delta to neighboring masses indicating a charge state >1 were deconvoluted to singly charged analogues. The scan was only further processed when the carbohydrate intensity was above the specified intensity threshold. The complete mass peak list was then subtracted from its (singly charged) parent ion mass. The thereby generated (negative) mass numbers (=‘parent offsets’) were collected in a separate matrix. By assuming a minimum peptide mass of 500Da (or roughly 5 amino acids), offset numbers <(500 – (parent mass)) were excluded. The generation of the ‘parent offset’ numbers was repeated for every scan containing a particular carbohydrate fragment. The collected ‘parent offset’ numbers (for a particular carbohydrate fragment) were finally trimmed to the specified mass range (0–2000Da, except otherwise specified), converted into absolute values and binned into 0.01Da windows, and visualized using the ‘histogram’ function.

Alternatively, the ‘parent ion offset’ binning of the complete shotgun proteomics dataset was performed using exactly the same approach as described for spectra filtered for the occurrence of certain carbohydrate fragment. Furthermore, the intensity normalized low mass bins for spectra containing a certain oligosaccharide chain were generated by binning the low mass range, after normalizing every mass peaks within a scan to the total peak intensity (100). Peaks with a relative abundance below 0.5 were not further considered. High-resolution mass binning to obtain the accurate mass of the oligosaccharide modification was performed by binning a focused mass range from 350–425, or 1150–1250m/z respectively, to achieve a resolution of 0.01 units bin size. Oligosaccharide variable modification masses for PEAKS database search were obtained by annotating the most abundant isotope within an oligosaccharide isotope cluster (after mass binning at a bin size of 0.05m/z, from 150–2000Da, and normalisation) using the ‘mspeaks’ function.

### (D) METABOLOMICS

#### Analysis of released nonulosonic acids and activated sugars

Nonulosonic acids were acid released from solid phase extraction purified peptide fractions (section B), DMB labelled and analysed using a slightly modified approach from Kleikamp et al., 2020.^[8a]^ Briefly, DMB labelled lysates were analysed by a Waters Acuity UPLC system using a C18 1.7μm BEH separation column coupled to a QE Focus Orbitrap mass spectrometer. The Orbitrap was operated in ES+ mode alternating full scans and small mass window fragmentation scans from 390-525Da. Mass windows were isolated at a window of 5.5Da and fragmentation was performed at a NCE of 26. Mass spectrometric raw data were analysed using the Matlab scripts as described in Kleikamp et al., and by using the Thermo Xcalibur Qual Browser. Activated sugars were analysed from fresh *Ca.* Kuenenia stuttgartiensis culture biomass. Sampling and metabolite extraction was performed according to the protocols described in Lawson et al., 2020.^[28]^ Briefly, cell broth was harvested, centrifuged in the cold and pellets were immediately dissolved and vortexed for 60 seconds using an extraction solvent precooled to −80°C (40/40/20 acetonitrile/methanol/water). Samples were further placed to - 80°C for 30 minutes and centrifuged (10.000 rpm, 4°C, 5min). 1ml of cell-free supernatant was collected and stored at −80°C for activated sugar analysis. Identification of activated sugars was performed according to Schatschneider et al.^[47]^ Briefly, LC-Orbitrap-MS analysis was performed using an ACQUITY UPLC M-Class liquid chromatography system coupled to a QE plus Orbitrap mass spectrometer (Thermo Fisher Scientific). Chromatographic separation was performed using a ZIC-pHILIC column (1.0×150 mm, 5μm, Merck, Germany) using 20mM ammonium carbonate in water, pH 9.1 as mobile phase A and 100% acetonitrile as mobile phase B. The flow rate was set to at 40μL/min. After 2.5 min constant at 82.5% solvent B, a gradient to 40% solvent B was performed over 15 minutes, followed by a gradient to 15% solvent B over 5 minutes. The mass spectrometer was operated in full scan mode acquiring signals from 150–700m/z in ESI negative mode at −2.5kV and 70K resolution with an AGC target of 3e6 and a max IT of 200ms. Raw MS data were investigated for activated sugar mass peaks using XCalibur 4.1 (Thermo Scientific, Germany).

### (E) GENOMICS

#### Metagenomic sequencing

DNA from the *Ca.* Brocadia sapporoensis enrichment culture was extracted using the DNeasy UltraClean Microbial Kit (Qiagen, The Netherlands). Following extraction, DNA was checked for quality by gel electrophorese and by using a Qubit 4 Fluorometer (Thermo Fisher Scientific, USA). Metagenomic sequencing was performed by Novogene Ltd. (Hongkong, China). Briefly, for library construction, a total amount of 1μg DNA per sample was used as input material. Sequencing libraries were generated using NEBNext^®^ Ultra™ DNA Library Prep Kit for Illumina (NEB, USA) following manufacturer’s recommendations. The DNA sample was fragmented by sonication to a size of 350bp, then DNA fragments were end-polished, A-tailed, and ligated with the full-length adaptor for Illumina sequencing with further PCR amplification. PCR products were purified (AMPure XP system) and libraries were analysed for their size distribution using an Agilent 2100 Bioanalyzer, and quantified using real-time PCR. The clustering of the index-coded samples was performed on a cBot Cluster Generation System according to the manufacturer’s instructions. After cluster generation, the library preparations were sequenced on an Illumina HiSeq platform and paired-end reads were generated. *Ca.* Kuenenia stuttgartiensis enrichment culture sampling and sequencing is described in Lawson et al., 2020.^[28]^

#### Metagenome assembly and binning

Raw reads were quality checked with FastQC v0.11.7 (http://www.bioinformatics.babraham.ac.uk/projects/fastqc/), and low-quality reads were trimmed using Trimmomatic v0.39^[48]^ with the default settings for pair-end reads. Trimmed reads were assembled for using metaSPAdes v3.13.0^[49]^ with default settings, resulting in 51,275 scaffolds of ≥1 kb. Metagenome binning was performed using three different binning algorithms: BusyBee Web^[50]^, MaxBin 2.0 v2.2.4^[51]^ and MetaBAT2 v2.12.1^[52]^. The three bin sets were supplied to DAS Tool v1.1.0^[53]^ for consensus binning to obtain the final optimized bins, which resulted in 47 metagenome assembled genomes (MAGs). Genome bins were assessed for completeness and contamination using CheckM v1.0.12^[54]^. As a result, 27 high-, 19 medium-, and 1 low-quality MAGs in accordance with minimum information about metagenome-assembled genome (MIMAG) standards^[55]^ were reconstructed. MAGs were classified taxonomically using GTDB-Tk v1.0.2 and the Genome Taxonomy Database (release 89). The reconstructed genomes were annotated through the NCBI Prokaryotic Genome Annotation Pipeline^[56]^. Annotation of the protein-coding genes was performed GhostKOALA tool^[57]^ (accessed October 2019) for Kyoto Encyclopedia of Genes and Genomes (KEGG) enzyme codes and supported with BLASTp (E value < 1e-20)^[58]^ searches against the NCBI non-redundant protein database.

#### Data availability

Raw sequencing data are available through the NCBI Sequence Read Archive (SRA) under accession number: SRR12344472. The MAGs are available at GenBank under accession numbers JACFMP000000000 to JACFOJ000000000. The BioProject accession number is PRJNA647942.

#### Phylogenetic analysis

Phylogenomic analysis of the Planctomycetes genomes was conducted using a concatenated alignment of 120 single-copy phylogenetic marker genes obtained using GTDB-Tk v1.0.2.^[59]^ A maximum-likelihood phylogenomic tree was inferred in IQ-TREE^[60]^ using the LG+F+I+G4 model recommended by ModelFinder^[61]^ with 10,000 Ultrafast bootstraps^[62]^. The tree was rooted using *Verrucomicrobium spinosum* DSM 4136 (GCF_000172155) as outgroup.

### (F) PROTEOME PHYSIOLOGY

#### Isoelectric point calculation

Protein isoelectric point prediction for the *Ca.* Kuenenia stuttgartiensis and *Ca.* Brocadia sapporoensis proteomes was performed using the isoelectric point calculation tool available through the sequence manipulation suite (SMS, https://www.bioinformatics.org), considering the pK values from EMBOSS.^[63]^ protein molecular weights were plotted against predicted isoelectric points using Matlabs ‘dscatter’ function. Protein amino acid statistics and net charge were determined using the ‘Prot pi’ tool (https://www.protpi.ch).

#### Protein sequence homology and conserved domain search

Protein homology and conserved domain search was performed against the NCBI non-redundant protein sequence database using blastp (protein-protein BLAST, https://blastuttgartiensisncbi.nlm.nih.gov) and TIGRFAMs using default settings.

